# Bacterial colony size growth estimation by deep learning

**DOI:** 10.1101/2023.04.25.538361

**Authors:** Sára Ágnes Nagy, László Makrai, István Csabai, Dóra Tőzsér, Géza Szita, Norbert Solymosi

**Author notes:** corresponding author(s): Norbert Solymosi.

## Abstract

The bacterial growth rate is important for pathogenicity and food safety. Therefore, the study of bacterial growth rate over time can provide important data from a medical and veterinary point of view. We trained convolutional neural networks (CNNs) on manually annotated solid medium cultures to detect bacterial colonies as accurately as possible. Predictions of bacterial colony size and growth rate were estimated from image sequences of independent *Staphylococcus aureus* cultures using trained CNNs. A simple linear model for control cultures with less than 150 colonies estimated that the mean growth rate was 60.3 *μm/h* for the first 24 h. Analyzing with a mixed effect model that also takes into account the effect of culture, smaller values of change in colony size were obtained (control: 51.0 *μm/h*, rifampicin pretreated: 36.5*μm/h*). An increase in the number of neighboring colonies clearly reduces the colony growth rate in the control group but less typically in the rifampicin-pretreated group. Based on our results, CNN-based bacterial colony detection and the subsequent analysis of bacterial colony growth dynamics might become an accurate and efficient tool for bacteriological work and research.

## Introduction

Bacteria reproduce by simple division, the rate of which is fundamentally influenced by the environment and the characteristics of the bacterium. The rate of bacterial multiplication is important for pathogenicity^1–3^ and food safety^4^. Therefore, the study of bacterial growth (multiplication rate) per unit of time can provide important data from a medical and veterinary point of view. Several developments for the automated monitoring of the growth rate exist. In liquid cultures, the quantification of turbidity (optical density), electrical conductivity, or redox potential can be used for this purpose.^5,6^ When cultured on solid media, the growth rate is estimated from the change in bacterial colony size. The most common solutions^7–11^ involve digital image analysis, used to detect colonies and then measure their size relying on a threshold-based approach. In comparison, sub-pixel correlation analysis in speckle imaging is a new frontier for this task.^12^

At the same time, the detection of bacterial colonies using neural networks is a promising approach^13,14^. However, when the aim is only to detect and count bacterial colonies, it is not necessary for the colony identifying feature (e.g. polygon, bounding box) to have the same size as the target colony. On the contrary, if we want to study growth rates, we need to be able to measure the size of the detected colonies as accurately as possible. This also means that we need to perform model selection according to such predictive measures of neural networks in order to obtain the most accurate results.

The aim of our work was to investigate the possibility of reproducing the bacterial colony growth results of Bärr et al.,^11^ using a convolutional neural network (CNN) previously trained on our annotated digital image dataset.

## Methods

To detect bacterial colonies in the Detectron2^15^ environment, 10 pre-trained Faster R-CNN models (R_50_C4_1x, R_50_C4_C4_3x, R_50_DC5_1x, R_50_DC5_3x, R_50_FPN_1x, R_50_FPN_3x, R_101_C4_C4_3x, R_101_DC5_3x, R_101_FPN_3x, X_101_32×8d_FPN_3x) were trained. For this purpose, our research group has previously created a manually annotated dataset (with bounding boxes enclosing the colonies). Since the images were of different sizes, they were transformed to a uniform size (6200*×* 6200 pt) for training. Each pre-trained model was trained through 100 epochs and validated after every 100 iterations. During validation, we always recorded weights with a smaller validation loss compared to the previous smallest weights. Thus, the training resulted in a collection of the best weights for each of the 10 pre-trained models.

We performed bacterial colony detection predictions using the best weights and an independent image collection. The dataset used to investigate bacterial colony growth consisted of the unannotated digital images generated and shared on Figshare by Bärr et al.^11^. The authors took digital images of 22 *Staphylococcus aureus* cultures in every 10 min. From each culture, 410 or 423 recordings were made. Of the 22 cultures, 8 were control (Ctrl), and 14 were pretreated with rifampicin (Rifa) for 24 h immediately prior to culturing (Table 1/A). Based on best weights, bacterial detection prediction was performed for images numbered 1-410. For the 410 records of 22 cultures (transformed to 6200 *×*6200 pt), predictions (bounding box coordinates, object classification probability) obtained with each model were stored. CNN training and predictions were performed on a Tesla V100 32GB GPU.

**Table 1.**
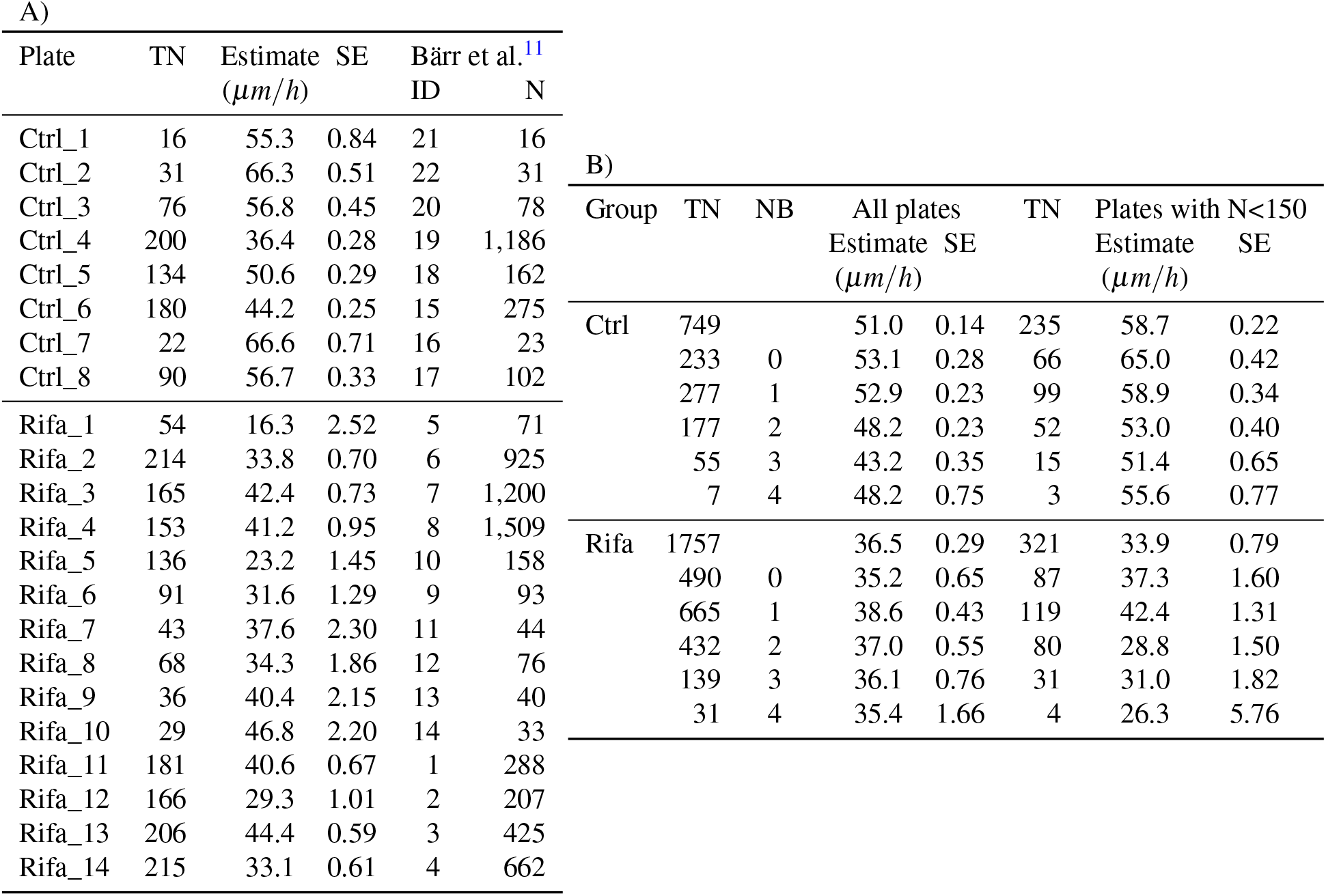
Hourly growth rate of the bacterial colony radius during the first 24 h. Table A shows values estimated by linear regression in each culture. The plate identifiers, taken from the name of the library of digital images hosted on Figshare by Bärr et al.,^11^ are the number of colonies tracked in the TN column, the ID column is the culture identifier used in the paper of Bärr et al.,^11^ and the N column is the number of colonies per culture. Table B shows the estimates of the radius growth within treatment groups using mixed-effect linear regression. Column NB indicates an additional grouping variable, the numbers indicate the number of close neighbors of colonies. The row without an NB value shows the estimates for all colonies.

Further processing of the data and plotting of the results were done in R-environment (v4.2.1)^16^ using the packages broom^17^, broom.mixed^18^, ggplot2^19^, sf^20^, tmap^21^, and xtable^22^. The bounding boxes around the colonies that were predicted with each model’s best weights were later filtered; only those with object prediction probabilities above 0.5 were retained for further analyses. Simple feature polygons were generated from their coordinates. From the predicted objects at 410 recording time, the ones with an object prediction probability greater than 0.95 were extracted. These represented the final state and size of the colonies. Based on our experience, we assumed that in the case of S. aureus, the colonies’ center does not shift significantly during growth. Therefore, when tracking the growth of colonies, we assumed that the bounding box describing the latest state of the colony mostly contains the area determined during the previous examinations. Accordingly, for each bounding box describing the final colony state, we extracted the bounding boxes predicted at the previous time points that fell completely within the final one. A series of these were used to estimate the growth rate of the colonies. The growth rate of bacterial colonies is influenced by how densely they are distributed in the culture. The number of other colonies in the surroundings of each colony that can affect its growth was characterized by the number of close neighbors (NB). We considered two colonies as close neighbors if the bounding boxes between them overlapped to any extent. In order to compare our results with the work of Bärr et al.^11^, we used the radius of the colonies to describe their size. We estimated the colony radius by taking half of the width of the predicted bounding boxes. The 10 models were evaluated by analyzing the series of the best weight predictions for all colonies. We selected the model that produced the largest number of elements in the series for each colony and the smallest radius based on the bounding boxes. Thus, the best model was created by training the pre-trained model X_101_32×8d_FPN_3x; therefore, the predictions obtained with this model are presented in further sections.

To estimate the appearance time of colonies for each culture, we extracted CNN’s first identified member of a colony growth series. For comparability with the Bärr et al.^11^ results, we used the per-plate linear model to estimate colony growth. Statistically, measurements on the same plate can be considered repeated measurements; therefore, we also used a mixed-effect linear model to estimate colony growth by treating the plate as a random factor. Following the work of Bärr et al.,^11^ we used the colony growth rates predicted during the first 24 h of culturing.

## Results

Regardless of the number of colonies, the trained neural network processed a single culture record in 0.31 seconds. This time included converting the image to a uniform size, predicting the bounding box of the colonies, and tabulating the data describing them. The bounding boxes on an arbitrary image of the Ctrl_1 culture are shown in Figure 1.

**Figure 1.**
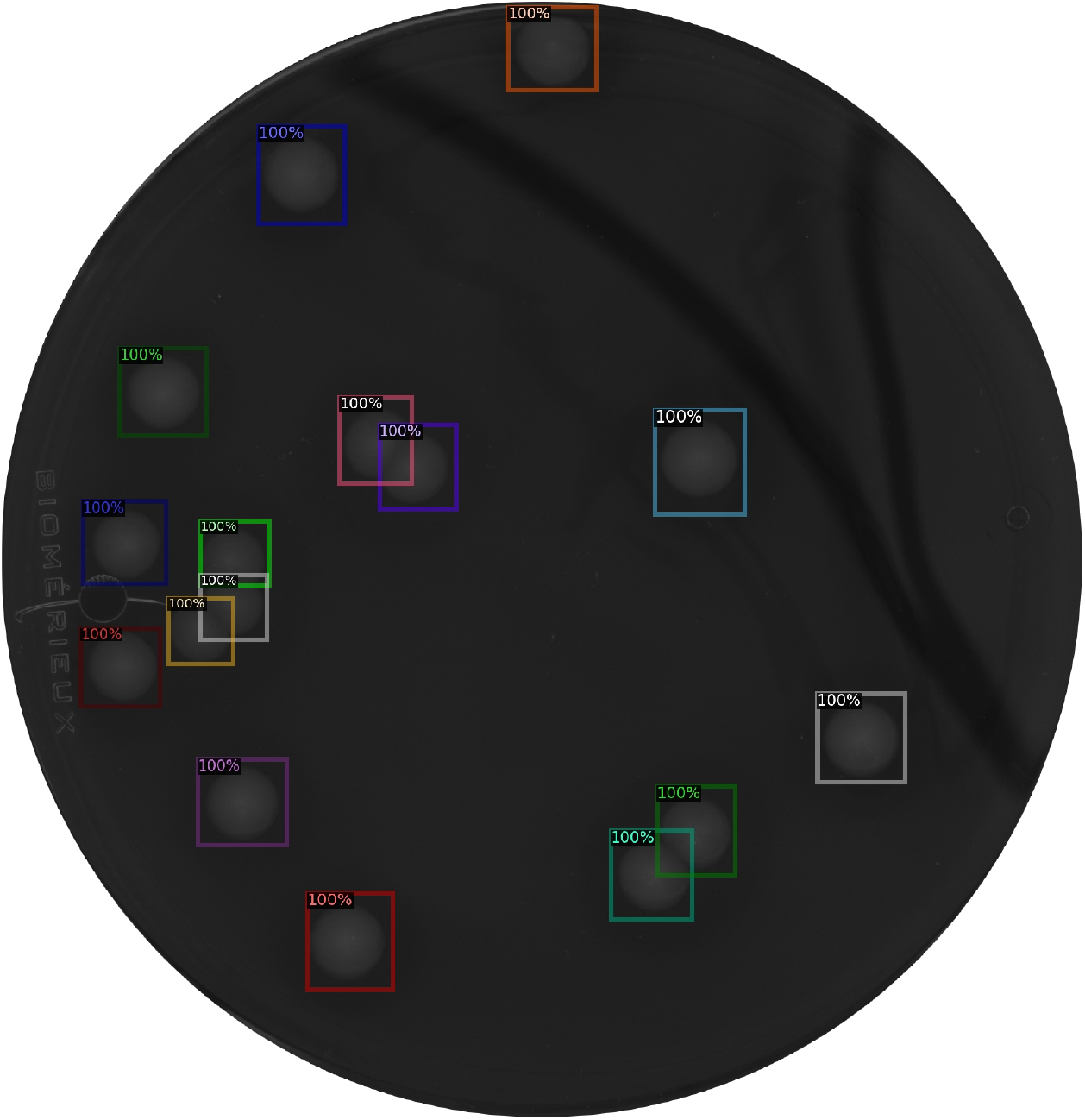
Bounding boxes on an image of Ctrl_1 culture predicted by the neural network. The colors have no meaning, they only aid separation in the case of overlapping boxes. The percentages indicate the confidence the algorithm assigned to the detected object

Fig 2 shows the time when the first colonies were detected in each culture. While the median time to first colony detection in the Ctrl group was 9.4 h, the median time to first colony detection in the Rifa group was 13.2 h, with a difference of 3.8 h.

**Figure 2.**
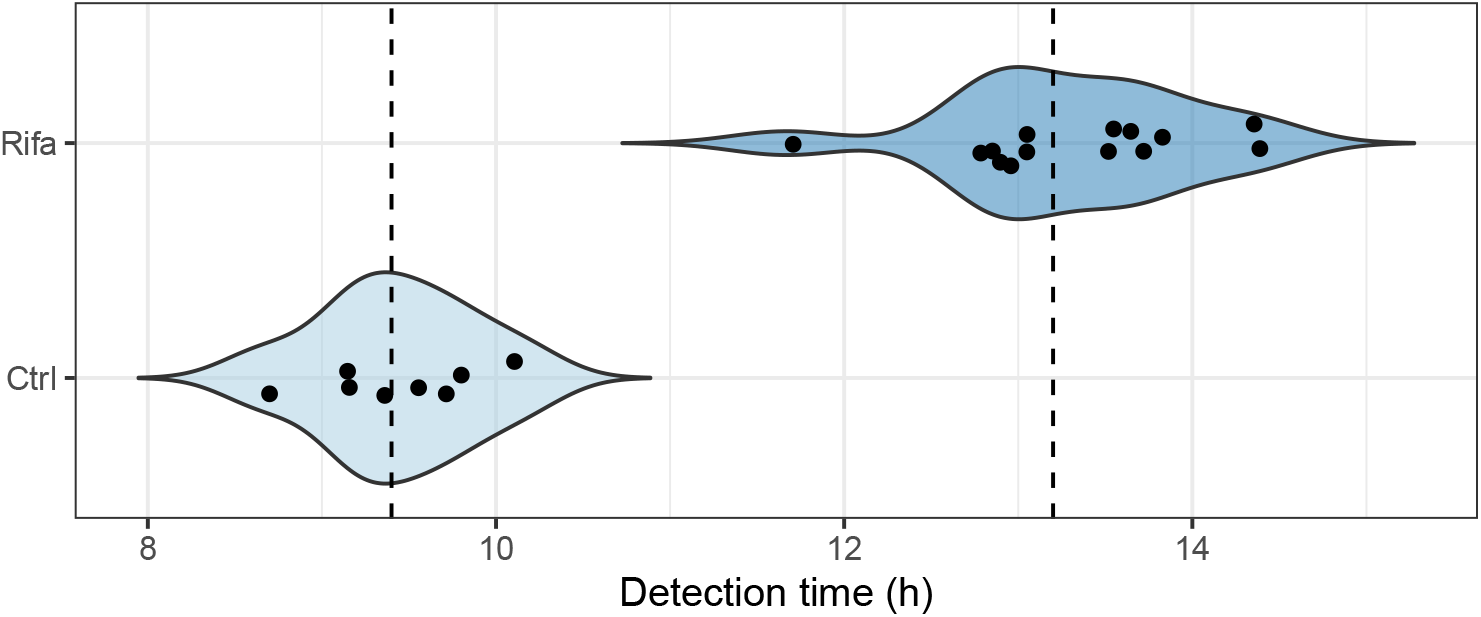
The time until the detection of the first colony in each culture in the two groups. Dashed lines indicate the median of the groups.

The growth curves of colonies are shown in Figure 3. Following the approach of Bärr et al.,^11^ we estimated the linear growth trend per culture for the first 24 h (Table 1). The average growth rate for Ctrl plates with less than 150 colonies was 60.3 *μm/h* (SD: 5.6). Using a mixed effect model, colony growth rate estimates for Ctrl and Rifa groups are summarized in Table 1.

**Figure 3.**
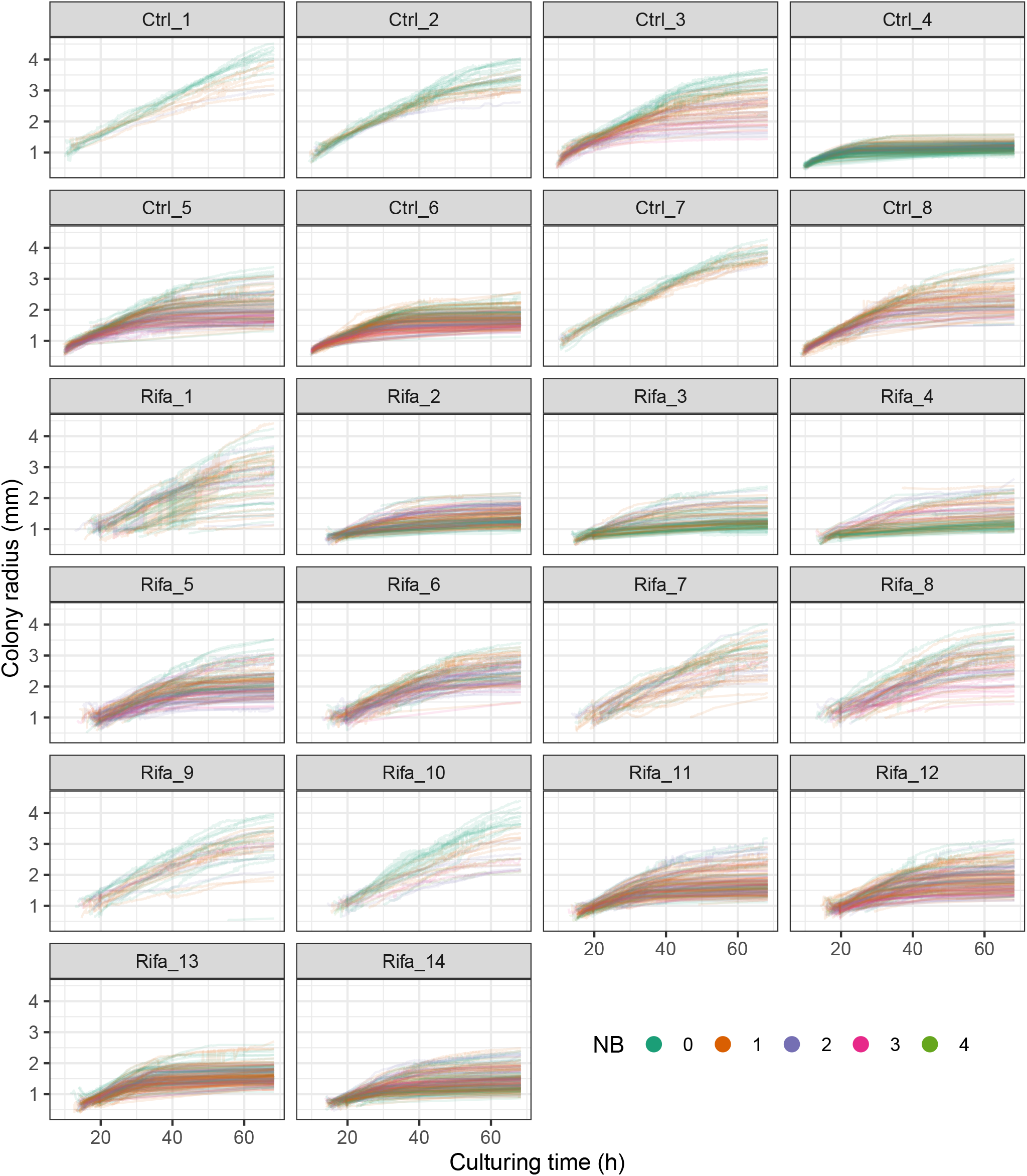
Growth curves of the colonies. Each curve describes the change in the radius size of a single colony (in Table 1, TN columns indicate their number). Their color indicates the number of close neighbors (NB) of the colony

## Discussion

The best model detected all colonies in the figure in 0.31 seconds, regardless of the number of colonies. Bärr et al.^11^ provided partial detection times for 16 cultures in their Supplementary Table 3, giving a mean per image of 8.8 s (SD: 7.35 s), which is 28 times our CNN result.

The median time to detect the first colonies in the Ctrl group (9.4 h) was 0.4 h later than that reported by Bärr et al.^11^ (9 h). In the Rifa group, however, instead of 17.4 h^11^, we obtained 13.2 h as the median of the appearance times. Thus, instead of 8.4 h^11^, we obtained a 3.8 h difference in the median of the appearance times of the first colonies in the two groups.

The hourly colony growth rates for the first 24 h in Ctrl group with less than 150 colonies were estimated by Bärr et al.^11^ to be 60.4 *μm*. Using the same statistical approach, our trained CNN estimated 60.3 *μm*, a difference of only 0.1 *μm* (1%). However, due to repeated measurements, we believe that a more correct approach is to use the mixed-effect model, which yields 58.7 *μm* for the same subset, with a 1.7 *μm* difference (2.8%) from the reference. For estimates that do not consider the number of neighbors, we can see that the rate for cultures with less than 150 colonies is always higher than the rate calculated from the sum of all cultures. This is more substantial in the Ctrl group (7.7 *μm/h*) and less in the Rifa group (2.6 *μm/h*). In both approaches, the Ctrl group shows that the growth rate decreases with the increasing number of neighbors up to the subgroup with 3 neighbors. Those with 4 neighbors show an increased growth rate, however, as Table 1 shows, the number of colonies with 4 neighbors is very low, therefore, the estimates for these are not really reliable. In the analyses using all Rifa cultures, we see that colonies without close neighbors grow at a lower rate than those with close neighbors, among which the increase in the number of close neighbors indicates a clear decrease in rate. No such regularity is seen in the Rifa cultures, with less than 150 colonies. Comparing Figure 3 showing the growth curves of colonies with the Supplementary Figure 10 of Bärr et al.,^11^ we see that the final colony sizes of our estimates exceed in several plates the values presented by Bärr et al.^11^ for the same ones. A visual inspection of the different cultures indicates that while the predicted bounding boxes are narrower in the case of the smaller colonies, more closely approximating the boundaries of the colonies, they can deviate significantly in the large colonies.

If the aforementioned minimal deviation in growth rates in the first 24 hours is reconsidered in this light, it can be explained by the fact that until the end of that period, the colonies are still quite small, and the predictions do not distort the size of the bounding box. We believe that the more inaccurate bounding box estimation of large colonies may be because the images used in the training set were taken from cultures that were incubated for 24-48 hours. As a consequence, only a few colonies could have grown as large as on the 68-hour cultures of Bärr et al.^11^ This imprecision could be reduced by using a training set that includes longer incubation time with larger colonies.

Based on our results, we believe that CNN-based bacterial colony detection and bacterial colony growth dynamics analyses could become an effective tool for bacteriological work and research.

## Declarations

### Ethics approval and consent to participate

Not applicable.

### Consent for publication

Not applicable.

### Availability of data and material

The datasets used and/or analyzed during the current study are available from the corresponding author upon reasonable request.

### Author contributions statement

NS takes responsibility for the data’s integrity and the data analysis’s accuracy. IC and NS conceived the concept of the study. NS and SÁN participated in the computing, statistical analysis, and drafting of the manuscript. DT, GS, IC, LM, NS, and SÁN carried out the manuscript’s critical revision for important intellectual content. All authors read and approved the final manuscript.

### Authors’ information

No relevant information can be provided by the authors that may aid the readers’ interpretation of the article.

## References

1. Anderson, J., Eftekhar, F., Aird, M. & Hammond, J. Role of bacterial growth rates in the epidemiology and pathogenesis of urinary infections in women. J. Clin. Microbiol. 10, 766–771 (1979).

2. Fisher, R. A., Gollan, B. & Helaine, S. Persistent bacterial infections and persister cells. Nat. Rev. Microbiol. 15, 453–464 (2017).

3. Bourchookarn, A., Paddock, C., Macaluso, K. & Bourchookarn, W. Association between growth rate and pathogenicity of spotted fever group Rickettsia. J Pure Appl Microbiol 16.

4. McMeekin, T. et al. Quantitative microbiology: a basis for food safety. Emerg. infectious diseases 3, 541 (1997).

5. Madrid, R. E., Felice, C. J. & Valentinuzzi, M. E. Automatic on-line analyser of microbial growth using simultaneous measurements of impedance and turbidity. Med. & biological engineering & computing 37, 789–793 (1999).

6. Lindqvist, R. Estimation of staphylococcus aureus growth parameters from turbidity data: characterization of strain variation and comparison of methods. Appl. environmental microbiology 72, 4862–4870 (2006).

7. Levin-Reisman, I. et al. Automated imaging with scanlag reveals previously undetectable bacterial growth phenotypes. Nat. methods 7, 737–739 (2010).

8. Levin-Reisman, I., Fridman, O. & Balaban, N. Q. Scanlag: high-throughput quantification of colony growth and lag time. J. visualized experiments: JoVE (2014).

9. Barr, D. A. et al. Serial image analysis of mycobacterium tuberculosis colony growth reveals a persistent subpopulation in sputum during treatment of pulmonary tb. Tuberculosis 98, 110–115 (2016).

10. Vulin, C., Leimer, N., Huemer, M., Ackermann, M. & Zinkernagel, A. S. Prolonged bacterial lag time results in small colony variants that represent a sub-population of persisters. Nat. communications 9, 4074 (2018).

11. Bär, J., Boumasmoud, M., Kouyos, R. D., Zinkernagel, A. S. & Vulin, C. Efficient microbial colony growth dynamics quantification with ColTapp, an automated image analysis application. Sci. Reports 10, 16084, DOI: 10.1038/s41598-020-72979-4 (2020).

12. Balmages, I. et al. Use of the speckle imaging sub-pixel correlation analysis in revealing a mechanism of microbial colony growth. Sci. Reports 13, 2613 (2023).

13. Majchrowska, S. et al. Agar a microbial colony dataset for deep learning detection. arXiv preprint arXiv:2108.01234 (2021).

14. Pawłowski, J., Majchrowska, S. & Golan, T. Generation of microbial colonies dataset with deep learning style transfer. Sci. Reports 12, 5212 (2022).

15. Yuxin Wu and Alexander Kirillov and Francisco Massa and Wan-Yen Lo and Ross Girshick. Detectron2. https://github.com/facebookresearch/detectron2 (2019).

16. R Core Team. R: A Language and Environment for Statistical Computing. R Foundation for Statistical Computing, Vienna, Austria (2022).

17. Robinson, D., Hayes, A. & Couch, S. broom: Convert Statistical Objects into Tidy Tibbles (2023). R package version 1.0.4.

18. Bolker, B. & Robinson, D. broom.mixed: Tidying Methods for Mixed Models (2022). R package version 0.2.9.4.

19. Wickham, H. ggplot2: Elegant Graphics for Data Analysis (Springer-Verlag New York, 2016).

20. Pebesma, E. Simple Features for R: Standardized Support for Spatial Vector Data. The R Journal 10, 439–446, DOI: 10.32614/RJ-2018-009 (2018).

21. Tennekes, M. tmap: Thematic maps in R. J. Stat. Softw. 84, 1–39, DOI: 10.18637/jss.v084.i06 (2018).

22. Dahl, D. B., Scott, D., Roosen, C., Magnusson, A. & Swinton, J. xtable: Export Tables to LaTeX or HTML (2019). R package version 1. 8–4.

